# A Pipeline for a Primate Projectome: mapping every individual myelinated axon across the whole brain

**DOI:** 10.1101/2023.10.23.563679

**Authors:** Gregg Wildenberg, Kevin Boergens, Viktor Nikitin, Alex Deriy, Francesco De Carlo, Vincent De Andrade, Xianghui Xiao, Narayan Kasthuri

**Affiliations:** Department of Neurobiology, The University of Chicago; Argonne National Laboratory, Biosciences Division, Lemont, IL, USA; Department of Physics, University of Illinois, Chicago; Argonne National Laboratory, Advanced Photon Source, Lemont, IL, USA; National Synchrotron Light Source II, Brookhaven National Laboratory, Upton, NY, USA

## Abstract

We developed a pipeline to detail the morphology and complete trajectory of every long distance projecting neuron in large volumes of the non-human primate brain - a projectome. A projectome would organize how the hundreds of brain regions in a primate brain communicate with each other and provide a comprehensive classification of long distance neurons. The reason such projectomes do not exist already is that there is no single imaging technique with the resolution and sampling depth capable of producing one in primates. Here we address that gap by combining new ways to cut brains with minimal tissue loss, nanoscale x-ray microscopy imaging that reveals every individual myelinated axon (MA) and the location, size, and morphology of every neuron, and algorithms for tracing individual myelinated axons for millimeters to centimeters, across tissue gaps.

## 2 Introduction

A fundamental goal of neuroscience is to determine the organizational logic of brains. Brains are obviously parcellated, and in some large brains like the non-human primate (i..e, *Rhesus macaque*) and human, there are likely hundreds of anatomically and functionally distinct brains regions [1, 2]. Decades of experiments have suggested that information moves across these brain regions in specific ways critical for brain functioning [3] and disruptions of these pathways results in a range of neurological disorders (e.g., multiple sclerosis, autism, epilepsy, schizophrenia, aphasia) [4, 5, 6, 7]. However, our understanding of the how brain regions communicate is biased, limited by existing technology. For example, we have good models of the averaged trajectory of thousands of long distance myelinated axons (MA) imaged with MRI [7, 8], or of the long distance targets of a small fraction («1%) of individual neurons, revealed with optical microscopy [9, 10, 11, 12, 13]. Such data is clearly valuable, for example see ‘Van Essen’ diagrams [3], but these techniques leave open questions about brain-wide organizational logic at the cellular level, many of which we suggest could be answered with a projectome: the detailed paths of every individual long distance neurons with its downstream targets, *relative to each other* [14]. A projectome would at minimum classify potentially every single long distance projection neuron in every brain region by its targets and the dendrite morphology. Moreover, a projectome would reveal every single long distance pathway in a single brain, including some that have never been characterized before, the *∼*strength of connection of every brain region on its target, and the numbers of projection neurons across different cortical and subcortial areas - essentially creating a Rosetta stone for correlating with function or genetic expression in the future. By having a *whole brain* projectome, rather than a sub-division (e.g., visual cortex), will uniquely address variability in the organizations of brains by, for instance, quantifying:

1. How often the trajectory of individual neurons match the trajectories of nerve tracts (e.g., do individual axons branch when nerves branch and vice versa?).
2. How much variability is there in the composition of projection neuron classes and numbers of projection neurons across cortical and subcortical regions?
3. Across animals, is variability in an individual projection to a particular brain region compensated by downstream pathways (i.e., if Region X has a smaller projection to Region Y in one animal, does the projection from region Y to Z increase to compensate?).
4. Variability in disease states, in response to the disease.

We argue that such principles are hard, if not impossible to reveal with either averaged or sparse sampling approaches. Finally, a hallmark of species with bigger brains, ranging from mice to humans, is variability in behavior and macro-connectivity (e.g., MRI) [15, 16, 17, 18] and thus we argue projectomes of large brains are well poised to address the fundamental question of how variability in long distance communication between brain regions can underlie variability in behavior.

Part of the reason such maps don’t already exist is that there is no single imaging technique with the requisite resolution, sampling depth, and speed to image entire primate projectomes. Magnetic resonance imaging (MRI) approaches are fast, span whole brains, can be done *in vivo* but lack resolutions to parse out long-range properties of individual neurons. Fluorescence approaches (e.g., light sheet [19], expansion [20]) can resolve individual axons only if a small fraction (<5-10%) of axons are labeled and in addition, fluorescence techniques often require genetic access for expressing fluorophores. Electron microscopy (EM) offers sufficient resolution to resolve individual myelinated axons, labels all neurons and is species independent, but current 3D electron microscopy approaches are limited to volumes substantially smaller than whole primate brains [21, 22].

One promising option is to use synchrotron source x-ray imaging, which we have in the past demonstrated can image large volumes of brains at sub-cellular resolution [23] and indeed can image entire mouse brains, *∼*1 cm^3^, at sub-micron resolution [24]. Synchrotron micro and nano ray computed tomogrpahy (µXCT, nXCT) have several advantages relative to standard optical microscopy and electron microscopy:

1. depth of penetration [23]
2. infinite vertical tiling [25]
3. speed of imaging *∼*10 MHz for µXCT [23] and *∼*100 MHz for nXCT [26]
4. automated imaging [25]
5. compatibly with post-hoc EM [24]

However, imaging an entire macaque brain (*∼* 5 cm^3^) with x-rays poses at least 2 challenges. First, heavy metals used to stain brains for x-ray imaging (e.g., osmium, uranyl acetate, lead citrate) and epoxy resins need to diffuse through the entire brain. While it is now possible to stain entire mouse brains with osmium [27, 28, 29], the *∼*200 fold greater volume of macaque brains will still pose problems for chemical access primarily thru diffusion. One possibility is transcardial perfusion of stains [30], but such approaches remain unreliable, hazardous, and thus, unlikely to be used on scarce and valuable primate tissue. A second obstacle is that, even with the minimal scattering afforded by using x-rays, penetrating through multi-cm thick samples while maintaining focus and resolution remains problematic. A human projectome only exacerbate the problems by orders of magnitude.

Thus, we developed a pipeline that entails subdividing brains with minimal loss, imaging each sub-volume with synchrotron source x-rays (**Figure 1**) at resolutions sufficient to reveal the trajectory of every MA and the morphology of every cell, and algorithms to trace MA in subvolumes and across cut surfaces. While the work presented here is a proof-of-concept rather than a whole projectome, it solves many critical hurdles first necessary to surpass and lays the path forward for following most, if not all, individual MAs over large volumes of a primate brain.

**Figure 1.**
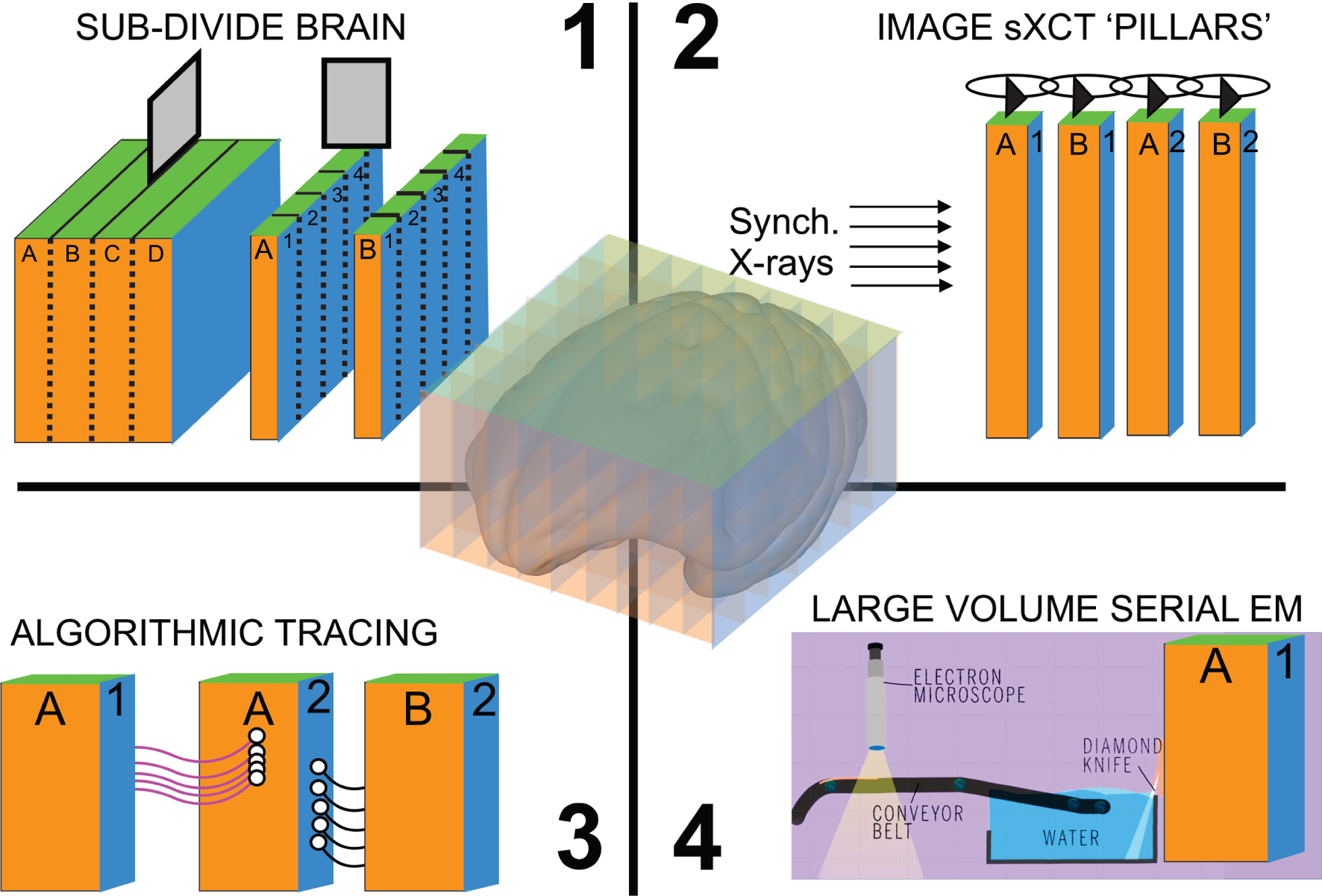
Proposed pipeline for x-ray primate projectome mapping. *Top left: 1*, a brain is cut into pillars by first cutting the brain into large flat slices (e.g., A, B, C, D, etc..) which are then cut a second time into thinner pillars (e.g., A1,A2,A3,A4 and B1,B2,B3,B4, etc…). *Top right: 2*, Each pillar, after being stained with heavy metals and embedded in resin, is imaged by synchrontron-source x-ray computed tomography (sXCT). *Bottom left: 3*, Individal MAs (depicted as lines between each pillar) are traced within and across each pillar to reconstruct a full brain projectome. *Bottom right: 4*, After projectome mapping, the same pillars are compatible with large volume serial EM for focused connectomic reconstruction of synaptic circuitry. *Middle*: Visual representation of primate brain sub-divided into a series of pillars.

## 3 Results

### 3.1 Cutting Brains with Minimal Loss

We first investigated whether a commercial system called the ‘Compresstome’, a new microtome that compresses tissue as it’s cut, developed for decreasing damage during cutting of living brain slices[31], could be used to tracing individual MAs across tissue sections. The Compresstome was recently used to track sparsely labeled fluorescent axons across serial macaque brain slices [9] but because this approach relied on labeling sparsity to follow axons across cut surfaces, the surfaces of the cut sections were not evaluated nor was the accuracy in following individual MAs across cut surfaces. We cut 300 µm thick serial slices of cortex brain tissue and stained each slice with Fluoromyelin, a fluorescent dye that labels individual MAs [32] (**Figure 2A**). We imaged opposing cut surfaces using confocal microscopy and found that we could easily identify many individual MAs across the cut surface, whether such axons were cut in cross-section, appearing as circles, or broadly parallel to the cut surface) (**Figure 2B**, red box and green circle, respectively). We next compared the relative surface integrity of a standard vibratome and compresstome brain section with scanning electron microscopy (SEM). Consistent with our ability to trace MA across cut surfaces, we found that the surface of compresstome sections were relatively preserved, with biological features like soma, nuclei, and individual MAs where reliably identifiable. Vibratome section surfaces, however, appeared uniformly jagged without any discernible biological features (**Supplementary Figure 1**).

**Figure 2.**
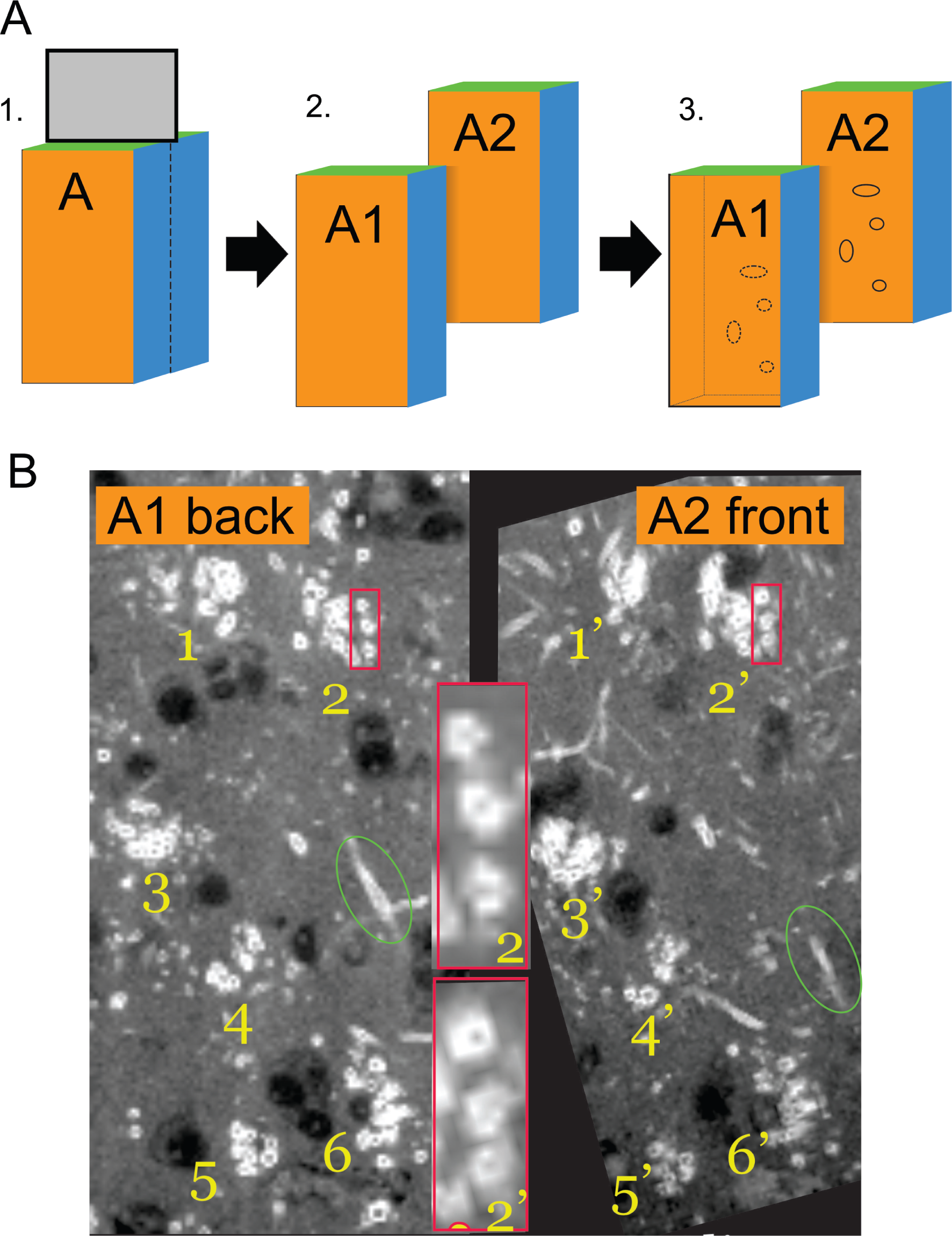
Individual MAs can be identified across brain sections cut with a Compresstome. **A.** Schematic of experimental design where a piece of tissue (A) is cut in half with a compresstome to produce two sections, A1 and A2. MAs in sections A1 and A2 are fluorescently stained with Fluoromyelin, and the matched surfaces are imaged (i.e., back of A1 and front of A2) by 2-photon confocal microscopy. **B.** Matched MAs from the back of A1 and the front of A2 can be manually identified (e.g., A1, 1 matches A2, 1’, etc…). *red box and inset*: zoom of cluster 2 and 2’ from A1 back and A2 front, respectively show three individual MAs that traverse the two sections. *Green circle*: an individual MA running in *∼*plane with the cut surface. Dark circles are negative contrast from unstained soma.

### 3.2 µXCT tomography of large brain pillars

We next used the Compresstome to cut a 2×4 pillar array of macaque cortex with each pillar *∼*0.7 mm × 0.5 mm × 8 mm in dimension, spanning across primary and secondary visual cortices (V1, V2 respectively) and the white matter tracts between (**Supplementary Figure 2**). Each pillar was then stained and embedded in a hard plastic resin using established EM protocols[33] which provide high contrast and physical stability for µXCT imaging[23, 24, 26] (**Figure 3A**). We developed custom cylindrical molds that reduce the sharp edges of the resin to minimize x-ray scattering[34, 35, 36] and that fit into a single field of view (FOV). Adjacent pillars were imaged using µXCT at *∼*600 nm^3^ resolution, and acquisition took approximately 75 min to image 1.3 x 10^10^ voxels (i.e., 2.9 MHz) (see Methods for how to access full data sets). In both V1 and V2 grey matter, individual neurons and their primary dendrites are clearly visible, and in white matter, numerous individual MAs can be resolved (**Figure 3B**). Additionally, manual inspection showed large fiducials (e.g., blood vessels) that cross two of the pillars, allowing identification of adjacent cut surfaces (**Figure 3C, Supplementary Movie 1.**).

**Figure 3.**
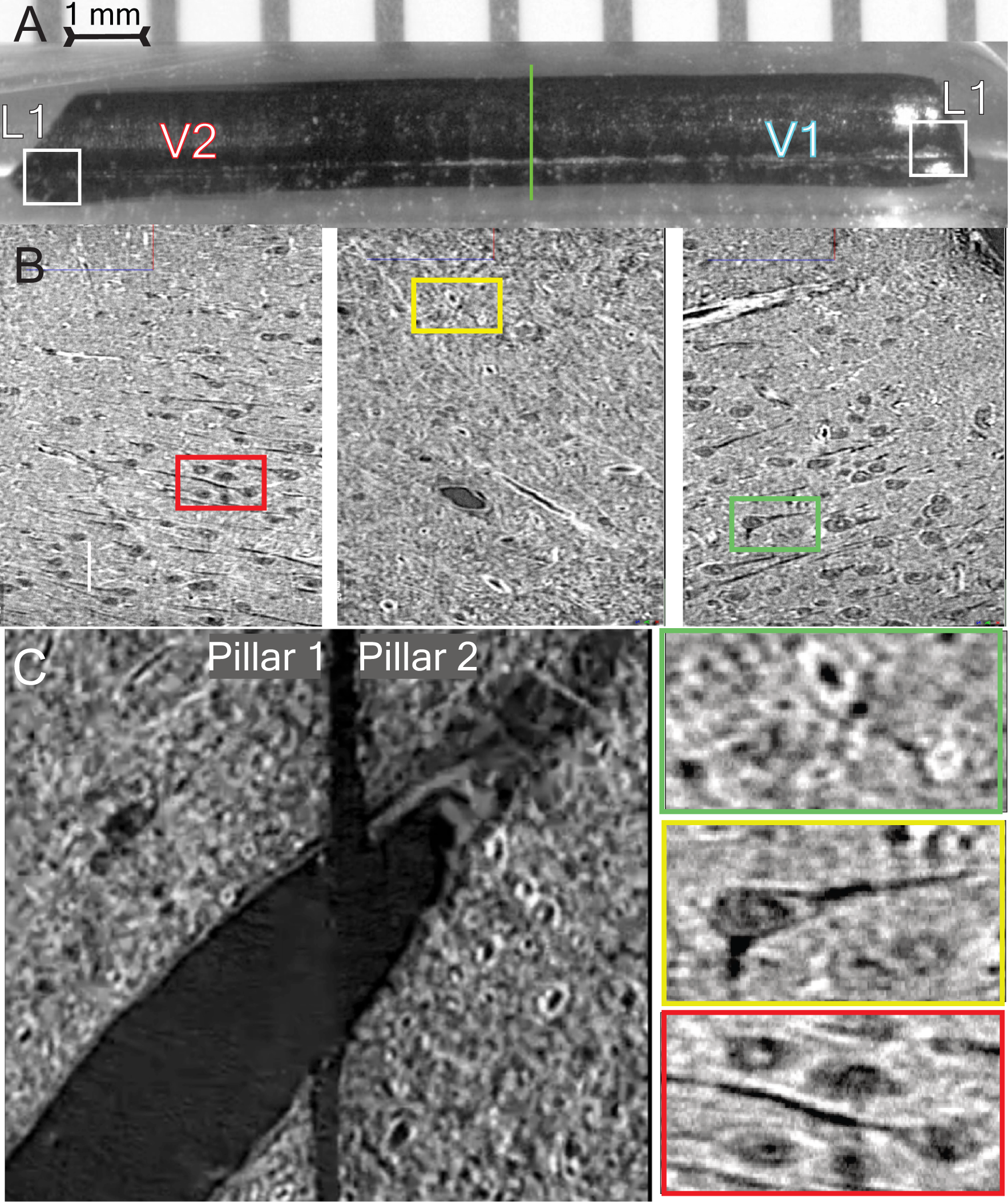
Pillar for sXCT imaging. **A**. Shown is an individual pillar (0.7 mm × 0.5 mm × 8 mm) spanning V1 and V2 with intervening white matter (green line) is prepared using the compresstome,stained for x-rays and embedded in a cylindrical resin mold prior to micron scale x-ray imaging. **B.** Sub-cellular features cab be readily identified including dendrites (*left and middle, red and green box*, and individual MAs (*right, yellow box*. *Below* 4x Zoom-ins from each example from corresponding colored boxes can be seen throughout the volume. **C.** Adjoining surfaces of two opposing pillars (1 and 2) are shown with a large blood vessel traversing across the gap. Scale Bar = 1 mm (A), 10 µm(B), 4 μm(C).

### 3.3 Tracing individual MAs across µXCT pillars and estimating tissue loss

We next used the µXCT datasets to ask whether individual MAs could be traced within and across across the gap between two pillars, and to calculate the tissue loss between pillars. We traced 31 readily identified MAs that traversed the two pillars (**Figure 4, and Supplementary Figure 3**)and we were able to follow 30 MAs across the entire 700 µm plane of the two pillars, and the one that could not be fully traced de-myelinated within the volume of pillar 1 but could be fully traced through pillar 2. The average length of MA traced was 1.4 mm, and in total we traced 86.99 mm of MA across the two pillars. We then used these reconstructed MAs to estimate the tissue loss, or “kerf”, between two pillars. Briefly, we fitted lines to each MA reconstruction that traverses the gap between the two pillars and estimated the distance between the ends of the MA reconstructions at the edges of each pillar (see Methods). Using this method we averaged the kerf over all 31 MAs as 697 nm (i.e., the loss from cutting is about 1 pixel of µXCT data). Overall, we conclude that: 1) large pillars can be readily stained and automatically imaged by µXCT, 2) there is minimal tissue loss between pillars, and 3) MAs can be manually followed throughout and across pillars even with micron scale x-ray imaging.

**Figure 4.**
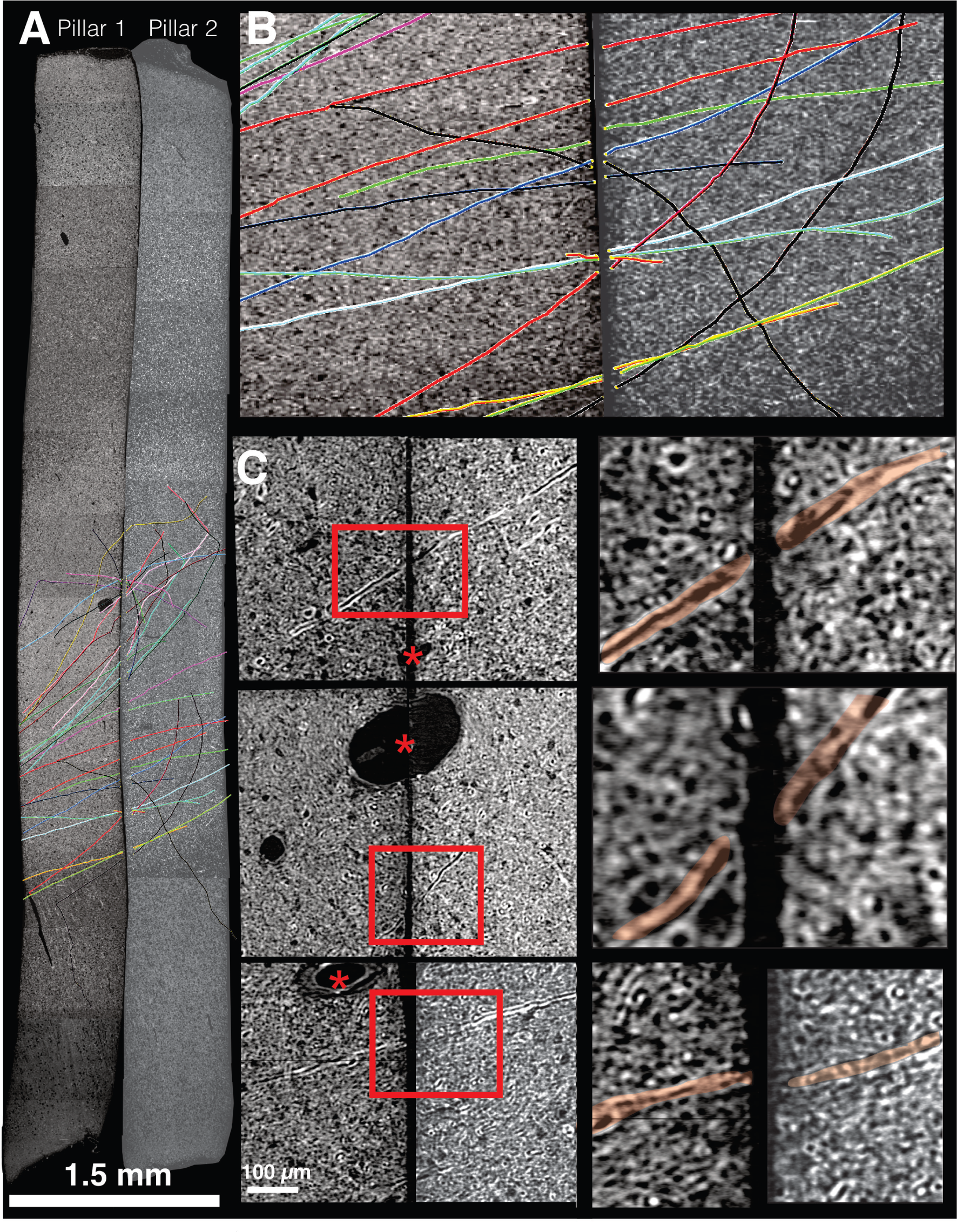
Individual myelinated axons can be traced across µXCT pillars. **A.** 2D image showing the entire length of pillars 1 and 2 overlayed with manual reconstructions of individual MAs that traverse the two datasets. Each colored line refers to an individual MA. **B.** Zoom-in of A on region where individual MAs were traced across pillars 1 and 2. **C.** Zoom-ins of 3 different regions showing individual MAs traversing pillars 1 and 2. *Red boxes*: regions further zoomed-in and shwon on right side with traversing MAs highlighted in red. Red asterisks mark blood vessels that traverse the two pillars. Scale bars = 1.5 mm (A), 100 µm (C).

### 3.4 Synchrotron *nano*-x-ray tomography for tracing MA

Although µXCT has the advantage of accommodating very large samples, even up to whole mouse brains [24], we found the resolution of µXCT was insufficient for automated segmentation of MAs and joining MA segments across the gap, both which would be necessary to scale this pipeline to whole brains. Thus, we next investigated nano-xray tomography (nXCT) imaging of brains as a higher resolution option for projectomics. We collected multiple nXCT datasets at multiple resolutions, using different nano x-ray imaging approaches, and at different synchrotron beamlines. We ultimately created two datasets, a 30 nm resolution, (*∼*50 µm^3^) volume from white matter in primates acquired on a Transmission x-ray Microscope (TXM) at the Full-field x-ray Imaging beam-line of the National Synchrotron Light Source II at Brookhaven National Laboratory (BNL), and a 100 nm (*∼*250 µm)^3^ volume of lower layers (L6) of mouse cortex at The European Synchrotron Radiation Facility (ESRF) using holotomography[37, 26, 38].

With 30 nm isotropic voxel resolution, we could reliably trace 100 percent of all MA within the volume, in dense white matter and across de-myleination events like nodes of Ranvier. We first asked how well such resolution could support tracing across kerfs. We produced computational gaps by removing slices from complete white matter nXCT stacks to mimic kerfs of different sizes (**Figure 5A**) to determine how accurately humans could trace MAs across different sized divides (See Methods). We removed 1.5 and 3 µm kerfs, *∼*2x and 4x larger than the actual gap we calculated for the compresstome sections, and had independent experts trace the same MAs across the 1.5 and 3 µm gap (**Figure 5B-C**). We found that experts agreed on the trajectories of 97% of the axons across a 1.5 µm gap, and that accuracy drops to 60% when the gap is 3 µm wide (104 MA fragments, mean length *∼*32 µm, total MA length 3.3 mm). Although 30 nm nXCT provides superior resolution, such imaging is currently limited to volumes of *∼*50 µm^3^ and would necessitate cutting 10,000 s of approx 1 mm diameter pillars to map even a 0.01% of macaque primate brain. Thus, we asked about mapping projectomes in a larger volume but coarser resolution nanoholotomography x-ray dataset [37] which we generated at ESRF at 100 nm^3^ voxel resolution. We first verified that humans could reliably trace individual MA throughout the volume (127 accurately axons traced; data not shown, see Methods). Furthermore, we found that at 100 nm voxel resolution, dendritic arbors could be clearly traced, and demyelination events like Nodes of Ranvier could be clearly visualized (**Figure 6A, Supplementary Figure 4**).

**Figure 5.**
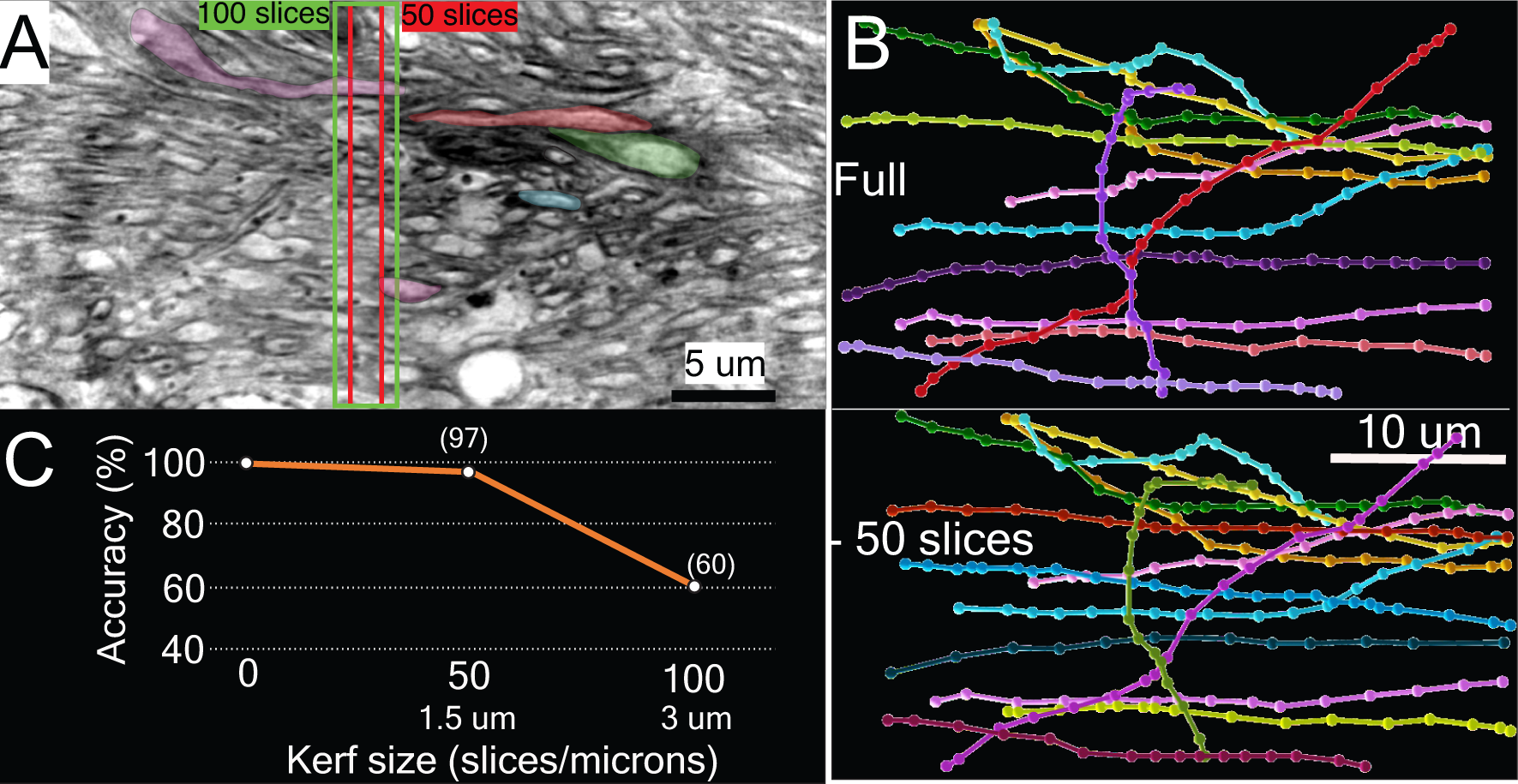
Individual MAs can be accurately traced across a computational kerf up to 1 µm. **A**. Schematic of experiment: the same 104 individual MAs were manually traced by three separate people in the same nXCT that is either: 1) full, 2) minus 50 slices (1.5 µm gap), and 3) minus 100 images (3 µm gap). **B.** Manual reconstructions of the same set of MAs from the full nXCT dataset (top) or the nXCT dataset with a 1.5 µm gap. Each color corresponds to an individual MA and the matching colors from the full and 1.5 µm kerf are the same MAs across datasets. **C.** Dot plot of x: computational kerf distance (µm) versus y: accuracy of tracing across the computational kerf. 97% of MA 1.5 µm kerf and 60% of MA can be traced across a kerk of 3 µm. Mean length +/-STD: 31.6+/-7.7 µm, total length traced = 3.26 mm. Scale bars = 5 µm (A), 10 µm (B).

**Figure 6.**
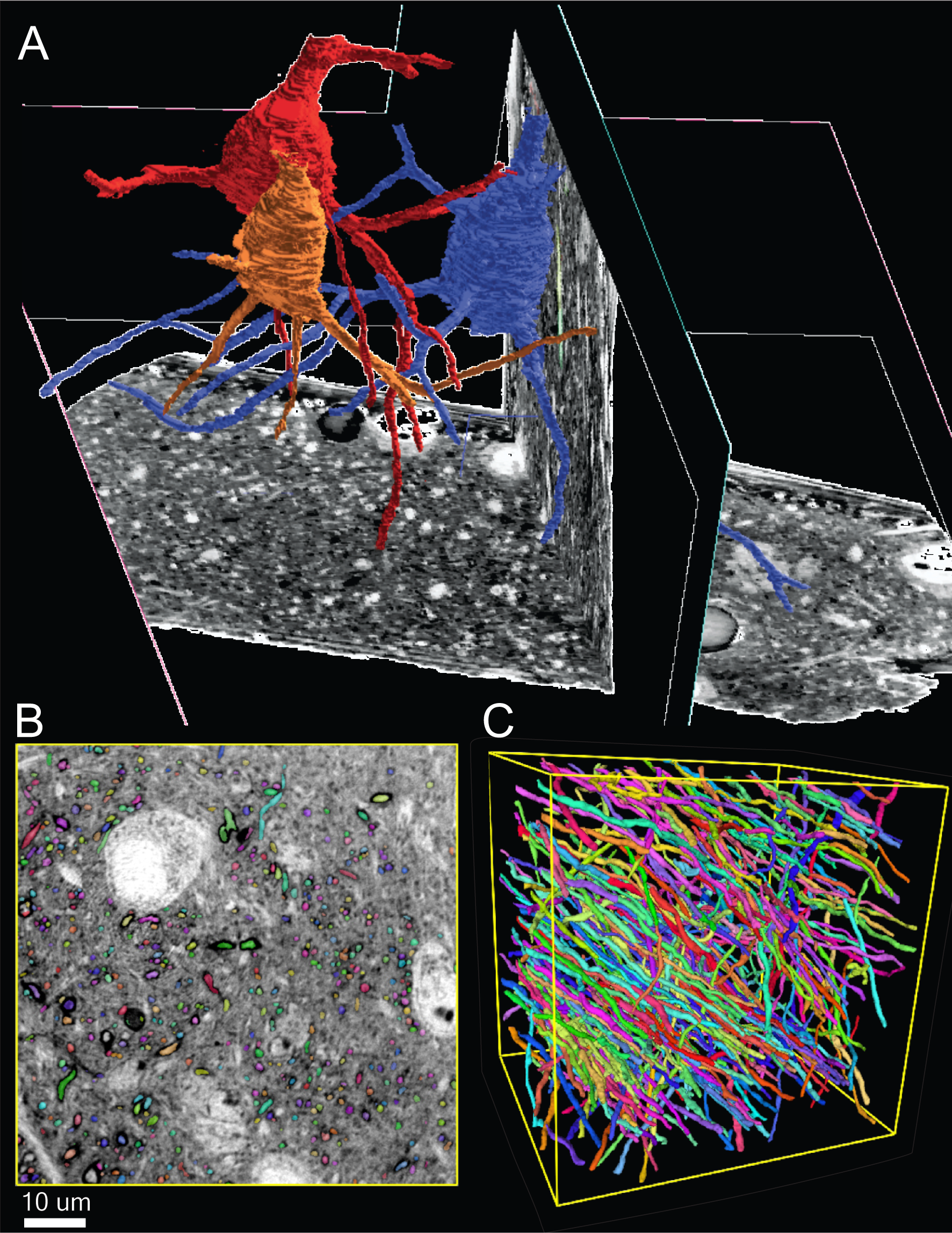
100 nm^3^ nXCT has sufficient resolution for reconstruction of neuronal morphologies and individual myelinated axons. **A**.3D renderings from manual reconstruction of three neurons overlayed on a 3D plane view of the raw nXCT data. **B**. Single 2D image from the nXCT volume overlayed with segmentation of individual MAs. **C**. 3D rendering of segmented MAs from (B). Scale bar = 10 (B).

We next asked whether algorithms used for electron microscopy data could reliably segment MAs from 100nm^3^ nXCT data. We found that a state of the art (SOA) EM segmentation algorithm, Local Shape Descriptors (LSD) [39], with minimal training could reliably identify nearly all MA profiles throughout a subvolume (116 µm^3^) of the nXCT data set (**Figure 6B**). We manually inspected 100 random MA segmentations and found 81% of MAs were accurately segmented throughout the entire volume (total length = 3.3 mm, average length = 35 µm) and all errors were splits (i.e., a single MA was labeled with 2 IDs). We found such splits were readily identified and fixed algorithmically: we first automatically identified putative split errors from the list of segmented objects by identifying tips of segments (i.e., where the segment ends). Second, we found MA tips that were within 250 nm and pointing at each other. We joined these ends and subsequent visual inspection found that 100% of theses repair were correct. Finally, we asked how well can algorithms trace MA across kerfs of 100nm^3^ nXCT data? To test this, we removed 19 sections, *∼*1.9 µm, from the segmented volume (*∼*2.5-fold greater than the estimated real gap of 697 nm), removed all unique identifiers, and used matching algorithms and directional predictions described above to reconstruct MA across gaps. We found that our algorithm performed 94.4% optimally in matching MA segmentations across the 1.9 µm divide **(Supplementary Figure 5)**. While such computational gaps are likely idealized (i.e., they do not contain cutting distortions or other artifacts), they suggest that tracing the vast majority of MA across Compresstome-generated gaps is feasible. Thus, with minimal training, available algorithms for automatic segmentation appear sufficient for our proposed pipeline for projectome mapping of nXCT data. Overall, these datasets confirmed several points that suggest 100 nm^3^ data appears to be a sufficient resolution for projectome mapping of individual MAs, and neuronal dendritic morphologies.

## 4 Discussion

Here we outline an experimental pipeline for mapping many to all individual myelinated axons and cell morphologies across large brains using synchrotron source x-ray imaging. We addressed several challenges:

1. methods for sectioning brains with minimal tissue loss and preparing for x-ray tomography.
2. identifying the minimal resolution needed to trace all individual MAs within volumes and across tissue gaps
3. automatic segmentation of individual MAs.
4. algorithms for joining segments of individual MAs across tissue gaps.

We found that with current x-rays resolutions (i.e., 100 nm voxels) we can trace MA within pillars with near 100% accuracy and across pillars with 97% human accuracy and 94% algorithmic accuracy across gaps we estimate are 2-fold wider than the actual gap between compresstome cut sections. Finally, we model the MA tracing success rate as P*≈*(Pseg*×*Pgap)^Ngap^, where Pseg is the fraction of axons within a pillar that are reconstructed without error, Pgap is the fraction of axons successfully traced across a kerf gap, and Ngap is the number of gaps jumped by the myelinated axon from end to end. Thus, the required success rate for an axon traversing on average 100 pillars (or 10 cm for 1 mm wide pillars) is Pseg*×*Pgap *≥* 97.8. Thus, we predict that algorithms could completely trace the long distance MA projections of 10% of all MA (10’s of thousands) in a cm^3^ volume (**Supplementary Figure 6**). This number could be increased with human intervention and error checking, which itself could be made more efficient by building expectations/priors for what MAs should look like using data collected from segmentation and error checking of MAs. These priors to identify possible sources of error (e.g., trifurcations, inappropriate loops, etc.) or to target human error checking could be scaled using the rising number of online tools to error checking the algorithmic output of connectomic data (EYEwire [40], FlyWire [41], and others [42, 43]). Second, there is now a public challenge for autosegmentation of MA through x-ray data: https://xpress.grand-challenge.org/ which will increase algorithmic efficiency.

### 4.1 A projectome of a primate hemi-brain

The projectome of an adult macaque hemi-brain, conservatively estimated at *∼* 5 x 3 x 5 cm, would create *∼*1500 pillars, 5 cm long and 1 mm wide as described above. These pillars could be imaged at existing synchrotrons, like the Advanced Photon Source (APS) at Argonne National Lab (ANL), using µXCT at 600 nm resolution in 78 days of continuous imaging (*∼* 50 MHz acquisition). Such data would allow for identifying every soma, tracing large fractions of dendritic arbors, tracing large caliber MA long distances, and detailing the full vasculature. A complete projectome, detailing the course of every MA in a volume will likely require 100 nm^3^ pixel size or better. We demonstrate such resolutions are currently achievable with nXCT and some beamlines have achieved such resolutions over a millimeter of penetration [38]. One critical factor for future projectomes is that beamlines are often upgraded, sometimes increasing their brightness by an order of magnitude. For example, the APS at ANL is currently upgrading with an expected increase of 500x brightness https://www.aps.anl.gov/APS-Upgrade. With the upgrade, µXCT at 600 nm^3^ of the hemi-brain would take 16 hrs and, assuming the major bottleneck in acquisition speed post-upgrade are detectors [44] and with the current availability of 6 GHz detectors, a 100 nm projectome would take 144 days.

## 5 Methods

### 5.1 Animal care and tissue processing

Perfusion procedures were followed according to animal regulations at the University of Chicago’s Animal Resources Center (ARC) and approved IACUC protocols. Rhesus macaques were housed in accordance with the Guide for the Care and Use of Laboratory Animals, Public Health Service Policy, and the Animal Welfare Act and Regulations, at the University of at Chicago, a fully AAALAC-accredited institution. All procedures involving animal care and use were approved by the IACUC at the University of Chicago.

### 5.2 Fluoromyelin staining

Tissue sections were provided by Precisionary Instruments, makers of the Compresstome. Paraformalde-hyde fixed zebrafinch brain was cut into 200 µm thick serial sections using the Compresstome, keeping track of the order of sections. Tissue was then stained with FluoroMyelin (Thermofisher Scientific, F34651) according to manufacture’s protocol. Stained sections were mounted with ProLong Gold Antifade Mountant (Thermofisher Scientific, P10144) and imaged with a Leica Sp5 Confocal microscope.

### 5.3 Tissue sectioning and staining

Primate brain tissue was prepared for EM as previously described [33]. Briefly, animals were deeply anesthetized until unresponsive. Primates were then transcardially perfused first with buffer (0.1M Sodium Cacodylate, pH 7.4) followed by fixative (0.1M Sodium Cacodylate, pH 7.4, 2% paraformaldehyde, and 2.5% glutaraldehyde). The brain was then extracted and postfixed for 24 hours in fixative at 4 *^◦^*C. A coronal piece of brain encompassing V1 was first cut out with a surgical scalpel followed by further excision of an area spanning V1 and V2 for either vibratome or compresstome sectioning. The visual cortex was identified using the calcarine and lunate sulcus, and a region just ventral to the lunate sulcus was selected that spanned V1 and V2 with intervening white matter. Tissue was then stained with heavy metals, dehydrated and embedded in plastic for electron microscopy as previously described [33].

### 5.4 Synchrotron x-ray computed tomography

Three x-ray datasets were generated for this study. First, the µXCT was produced at Argonne National Laboratory at station 2-BM using 20 keV energy, 10x objective and a detector size of 2448×2048. Each projection data volume was acquired at a 0.3 s exposure per projection with a total of 3000 angles totally 15 min/volume. About 5 scans were required to span the 8 mm axis at 0.345 µm pixel size, totaling about 75 minutes/pillar. Tomograms were reconstructed using publicly available scripts: https://github.com/tomography/tomopy-cli. Secondly, a nXCT data set was acquired with the transmission x-ray microscope (TXM) at the Full-field X-ray Imaging beamline of National Synchrotron Light Source II at Brookhaven National Laboratory [45, 46]. The x-ray energy used in the experiment was set to 8.5 keV. The projection images of the sample were recorded at 600 angles in 180*^◦^* with 0.3*^◦^* angle step. The exposure time for a single projection image was 0.1 s and the time for a tomography scan was 1 minute. Tomographic images of the sample were reconstructed from the projection images with Gridrec reconstruction algorithm [47] implemented in a Python package TomoPy [48]. The reconstructed tomography images have isotropic voxel size of 20.6 nm. The third dataset was collected at the European Synchrotron Radiation Facility (ESRF) (Grenoble, France) using holotomography and imaging parameters as previously cited[37, 26, 38].

### 5.5 Manual and automatic segmentation

MAs from µXCT data sets were manually reconstructed in Knossos (https://knossos.app/) using skeleton tracings. 3D reconstructions of neurons from the 100nm nXCT, and MA tracing across a computational kerf of the 30 nm nXCT data set were done in webknossos [49] ((https://webknossos.org/). Automatic segmentation of MA from the 100nm nXCT was done using the previously published local shape descriptor algorithm [39] which can be found here: https://github.com/funkelab/lsd.

### 5.6 µXCT gap estimation

To estimate the gap between the two adjacent pillars sectioned with a compresstome, 31 myelinated axons were identified that traversed the gap and skeleton-reconstructed using Knossos.Then we measured how strong the direction of the “edge” closest to the gap correlated with the edges as the skeleton tracing moved further away from the gap. To get a direction predictor of the skeleton as it enters the gap we used the last five edges closest to the gap and calculated the weighted average direction (according to the correlation coefficient). Next, we extended the skeleton across the hypothesized gap from both sides and varied the gap width to minimize the residual tension using the downhill simplex algorithm[50]. To measure the residual tension, we iterated over all axons, temporarily removed that axon and matched the axon stubs coming from pillar 1 onto the axon stubs coming from pillar 2 using the Clough-Tocher scheme[51] (because the relative alignment of the two pillars is not exactly known). We then used this interpolation to predict the position of the temporarily removed axon stub from pillar 1 and added the distance of its terminal to its partners terminal (from pillar 2) to the overall tension term.

### 5.7 Repairing MA segmentation splits

To repair splits we used the skeletons from manual annotations in the µXCT data sets that traversed pillars 1 and 2 as described in the methods section, µXCT gap estimation. We identified all endings of skeletons and measured the pairwise distance between all endings of all skeletons. We matched endings by stepwise increasing distance up to a cutoff of 250 nm and at every step matching endings that had a smaller distance. We rejected tentative ending pairings if the dot product of the two normalized direction vectors was larger than-sqrt(0.5). We produced an NML file detailing all matchings created this way and used webKnossos [49] to manually assess the performance of this matching.

### 5.8 Connecting MAs across a computational gap

To estimate how effectively we can reconstruct axons over a gap, we used the 100nm^3^ nXCT data and manually removed 19 slices (1.9 µm), creating two sub-datasets A and B. We then used the Kimimaro algorithm (https://zenodo.org/record/5539913) to skeletonize the segmentations derived automatically (**Figure 6B**). For each terminal on the surface of this artificial gap, we calculated the distance of each connected node along the skeleton (using the Dijkstra “shortest path” algorithm[52]) and measured average position of all connected nodes between 7 and 7.5 µm away (if none were, we substituted it with the position of the most distant node). We used the connecting vector from this average to the surface node as a direction predictor and extended the skeletons tentatively from both sides half-way into the gap. If this connecting vector was longer than 5 times the gap width it was rejected and a vector orthogonal to the cut surface was used instead. This lead to 374 skeletons coming from the A side of the artificial gap and terminating half-way into the gap and 383 skeletons coming from the B side of the artificial gap and terminating half-way into the gap. From here on, we considered these endings only in this half-way plane. We measured the pairwise distance of all endings from A and from B. We then used the Hungarian algorithm[53] to create a least-cost matching for these potential pairs, using the square root of the distance as the cost. We counted the number of correctly matched pairs and calculated the performance metric as a percentage of the highest theoretically achievable score.

## 5.9 Data availability

All datasets are freely available for download at BosssBD (https://bossdb.org/).

## Acknowledgements

We would like to sincerely thank Jian-Qiang for providing Compresstome sections. This research used the Full-field x-ray Imaging beamline 18-ID of the National Synchrotron Light Source II, a U.S. Department of Energy (DOE) Office of Science User Facility operated for the DOE Office of Science by Brookhaven National Laboratory under Contract No. DE-SC0012704

---

**Supplementary Figure 1.**
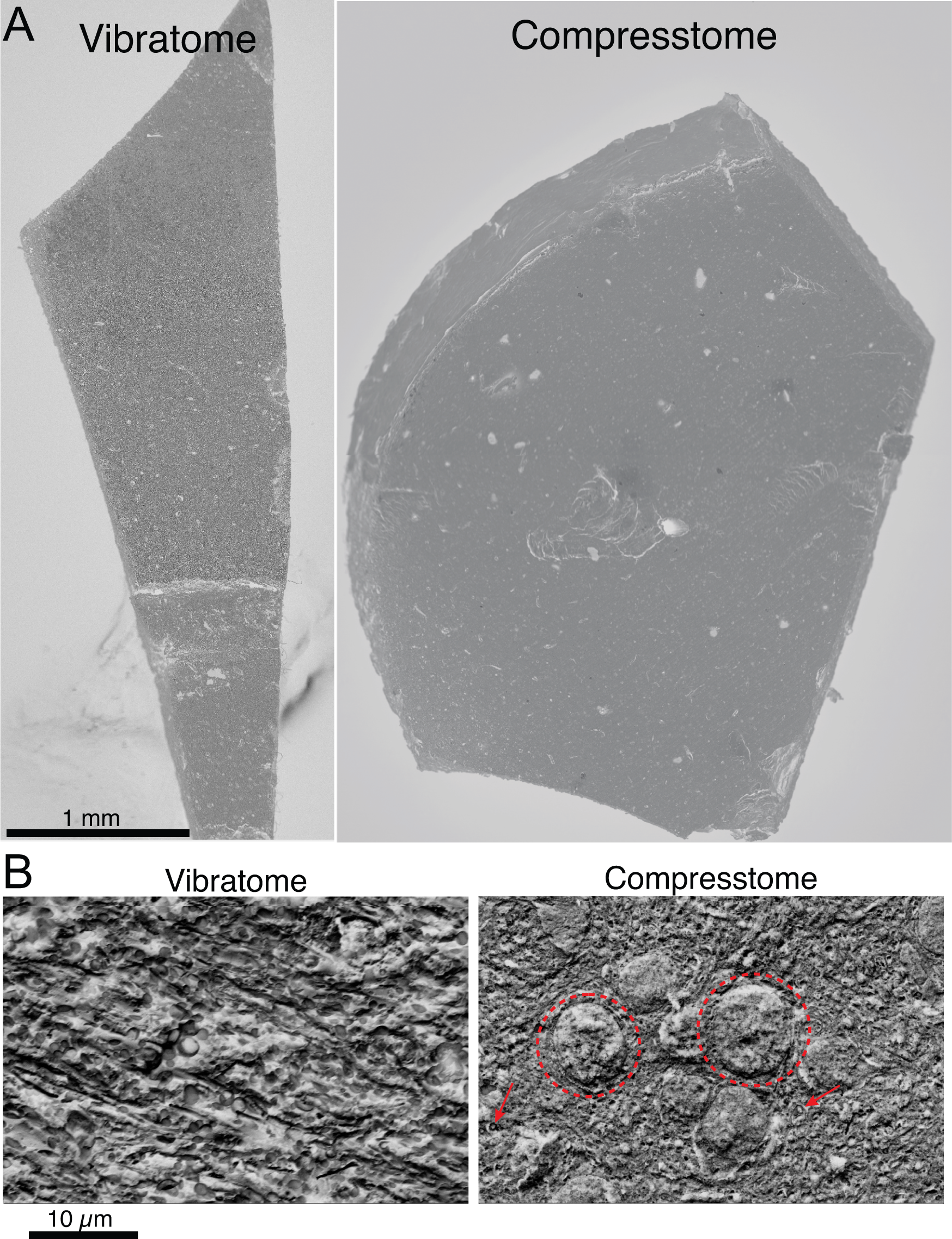
Compresstome sections retain biological architecture on very surface of the cut section. **A.** Low resolution SEM image of brain tissue cut with a vibratome (left) and compresstome (right). **B.** High resolution SEM image of the surface of a vibratome section (left) and compresstome section (right). Note that in the compresstome section, neuronal nuclei (dotted circle) and individual MAs (red arrows) can be readily visualized. Scale bar = 1 mm (A), 10 µm (B).

**Supplementary Figure 2.**
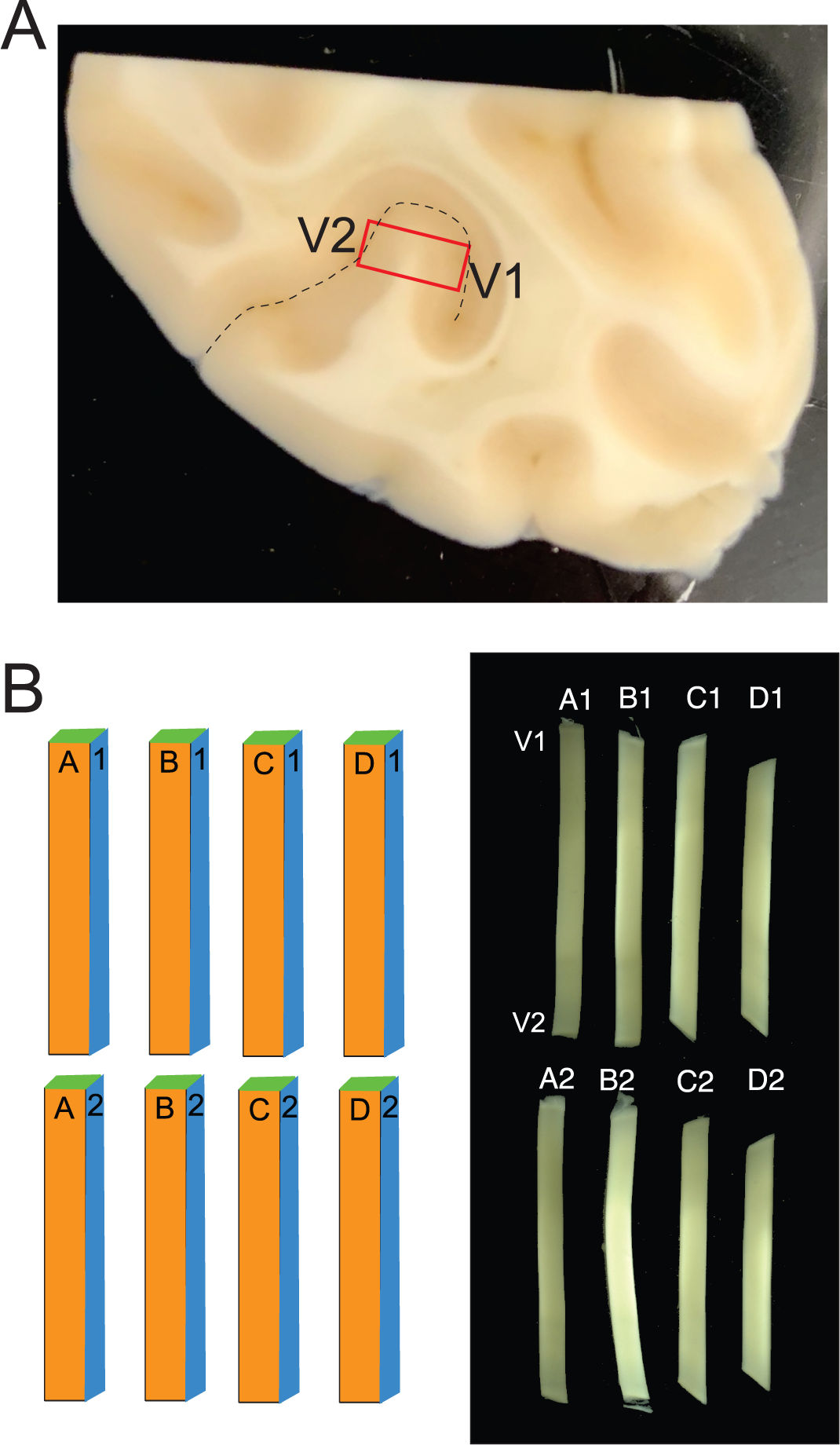
Brain region excised for primate projectome. **A**. A coronal piece of primate brain was cut using a scalpel and a small piece (red box) spanning V1 and V2 was further excised for Compresstome sectioning. V1 and V2 was identified by locating the calcarine sulcus (dashed line). **B**. Left: Cartoon of tissue pillar array made from Compresstome sectioning of V1/V2 brain tissue from (A). Right: bright field images of tissue pillar array spanning V1 and V2.

**Supplementary Figure 3.**
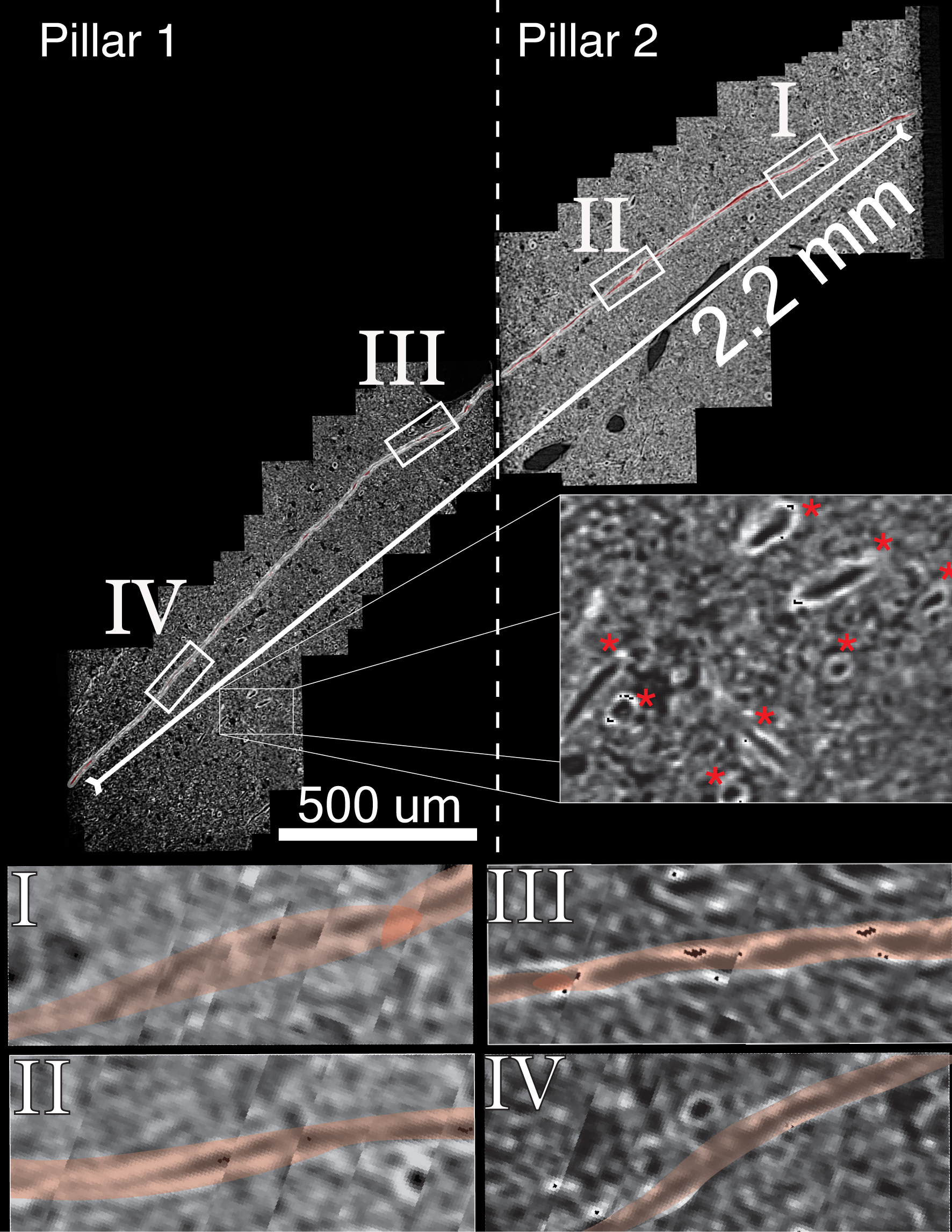
Montage of an individual MA traversing µXCT pillars 1 and 2. Individual images that follow a MA (outlined in red) across the two pillars were stitched together to show the 2.2 mm path and ability to be traced entirely. *Inset*: Zoom-in showing a cluster of additional MAs (red asterisks). Roman numerals I-IV correspond to zoomed in areas depicted below.

**Supplementary Figure 4.**
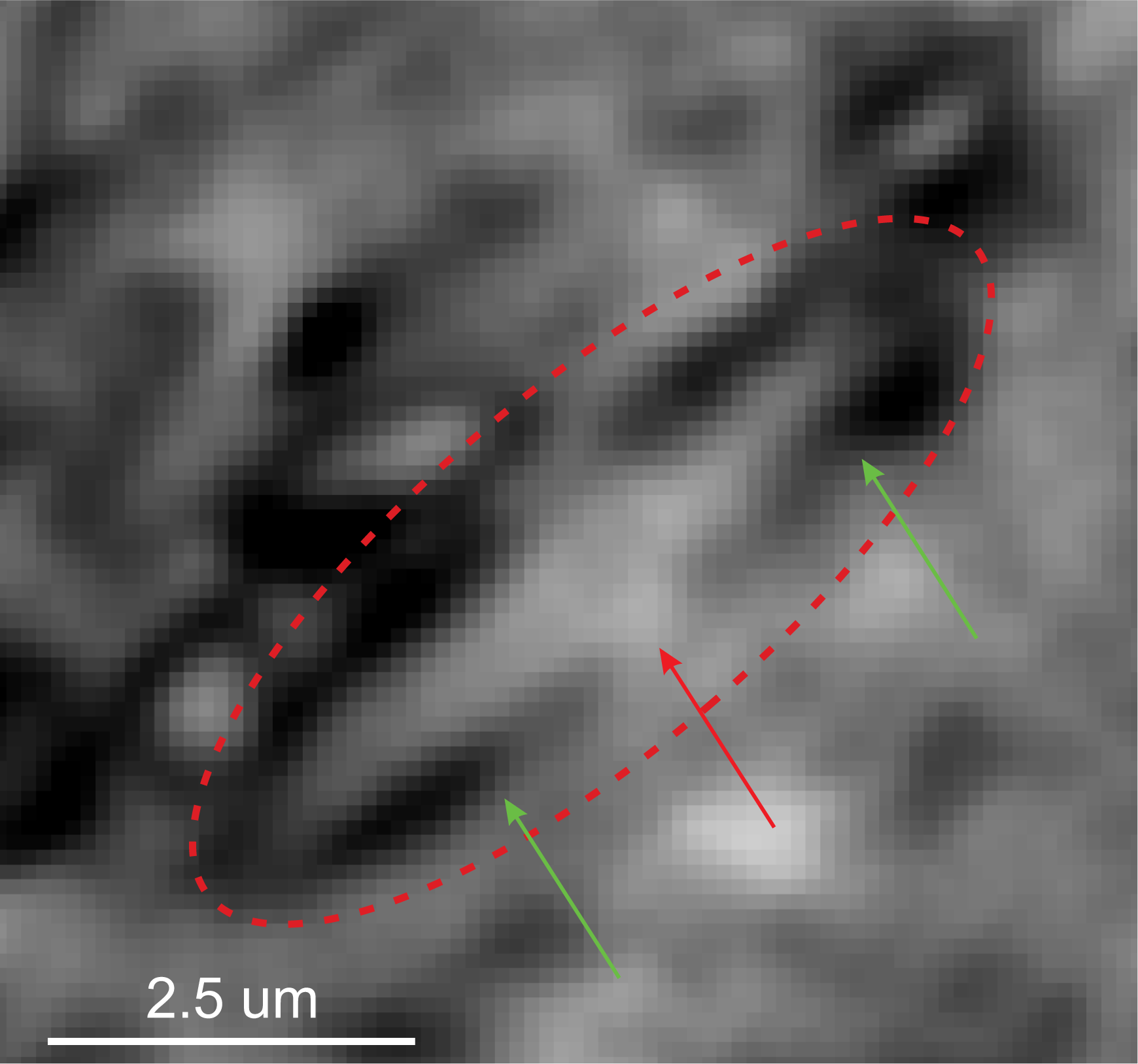
Nodes of Ranvier can be identified in nXCT. Zoom in of an individual myelinated axon (red dotted circle) with a Node of Ranvier. Red arrow points to the Node of Ranvier and green arrows point to the portion of myelin flanking the Node of Ranvier. Scale bar = 2.5 µm.

**Supplementary Figure 5.**
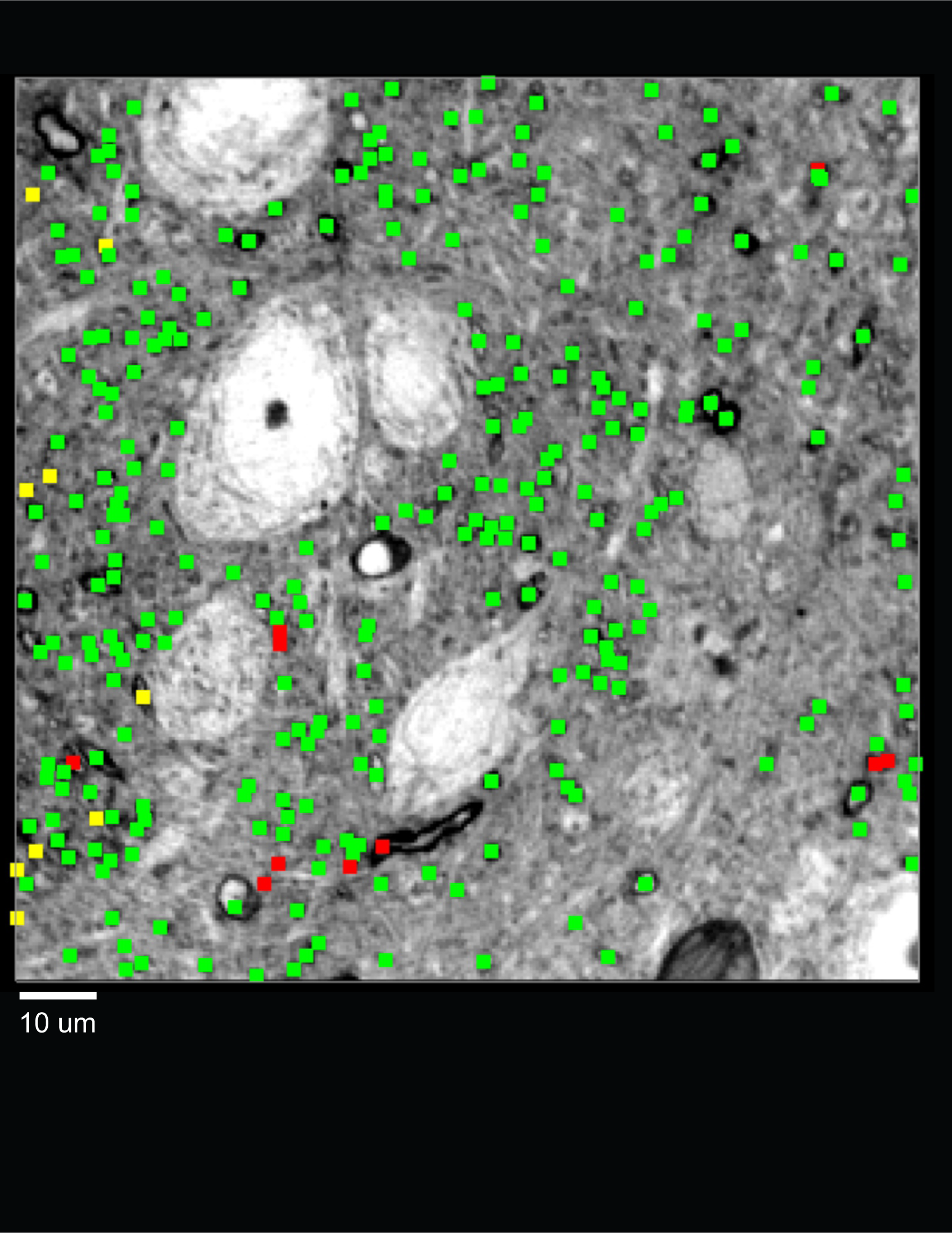
Individual MA segments can be joined across a computational gap. 100 nm resolution nXCT image showing the number of correct (green), incorrect (red) matches, and those that didn’t have a match across the gap (yellow) across a 1.9 µm gap. Scale bar = 10 µm.

**Supplementary Figure 6.**
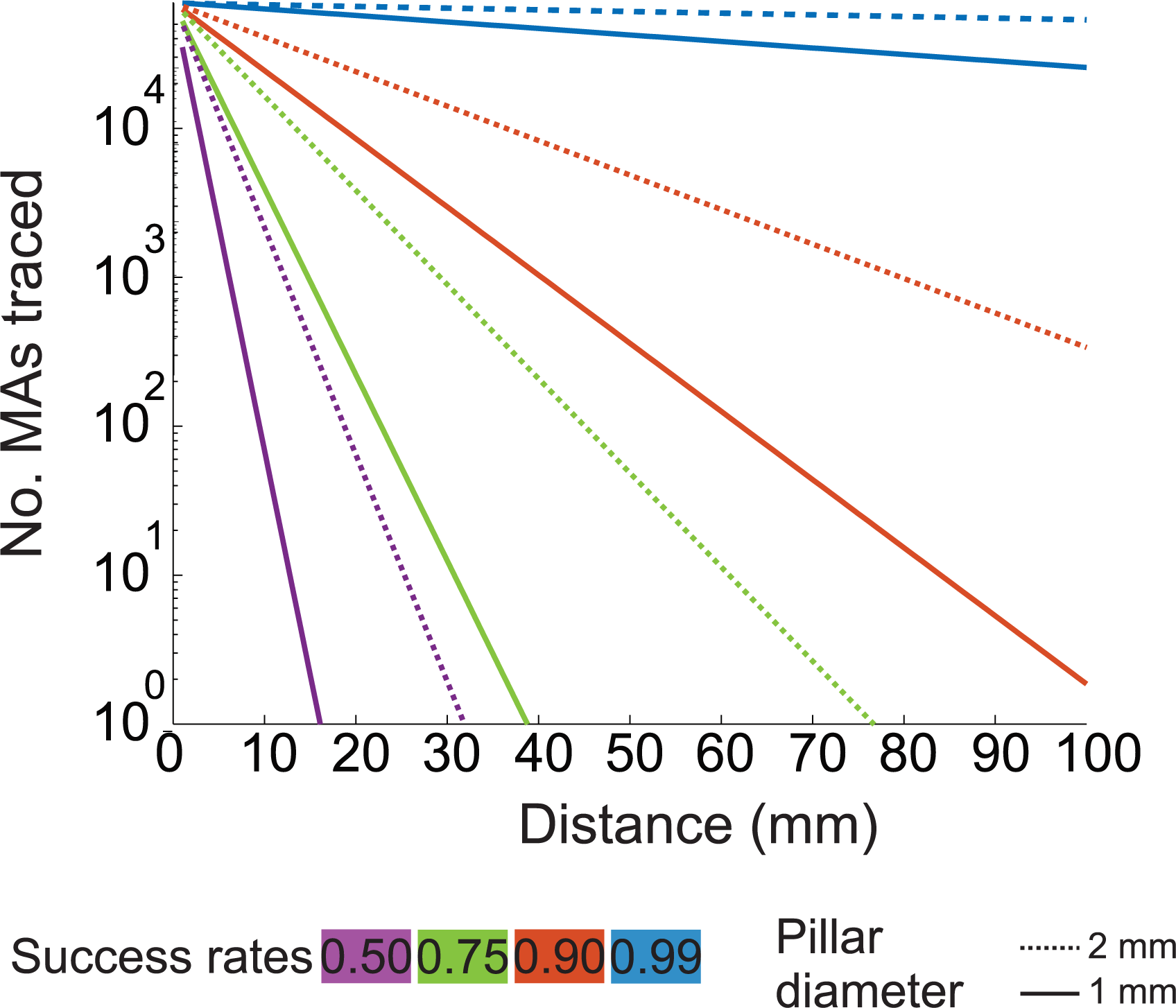
Estimated rates of successfully tracing myelinated axons across tissue pillars of different size. Plot of x: the number of neurons successfully traced versus y: the total distance neurons are accurately traced for different rates of successful tracing across pillars. Estimates were performed for both tissue pillars that are either 1 mm (solid line) or 2 mm (dashed line) diameter.

